# DNA transduction in *Sodalis* species: implications for the genetic modification of uncultured endosymbionts of insects

**DOI:** 10.1101/2020.12.02.408930

**Authors:** Chelsea M. Keller, Christopher G. Kendra, Roberto E. Bruna, David Craft, Mauricio H. Pontes

**Affiliations:** Department of Pathology and Laboratory Medicine, Pennsylvania State University College of Medicine, Hershey, PA 17033, USA; Department of Microbiology and Immunology, Pennsylvania State University College of Medicine, Hershey, PA 17033, USA

**Author notes:** Corresponding author: Mauricio H. Pontes, Penn State College of Medicine, Departments of Pathology, and Microbiology and Immunology, 500 University Drive, C6818A, Hershey, PA 17033.

## Abstract

Bacteriophages (phages) are ubiquitous in nature. These viruses play a number of central roles in microbial ecology and evolution by, for instance, promoting horizontal gene transfer (HGT) among bacterial species. The ability of phages to mediate HGT through transduction has been widely exploited as an experimental tool for the genetic study of bacteria. As such, bacteriophage P1 represents a prototypical generalized transducing phage with a broad host range that has been extensively employed in the genetic manipulation of *Escherichia coli* and a number of other model bacterial species. Here we demonstrate that P1 is capable of infecting, lysogenizing and promoting transduction in members of the bacterial genus *Sodalis*, including the maternally inherited insect endosymbiont *Sodalis glossinidius*. While establishing new tools for the genetic study of these bacterial species, our results suggest that P1 may be used to deliver DNA to many Gram negative endosymbionts in their insect host, thereby circumventing a culturing requirement to genetically manipulate these organisms.

**Summary:** A large number of economically important insects maintain intimate associations with maternally inherited endosymbiotic bacteria. Due to the inherit nature of these associations, insect endosymbionts cannot be usually isolated in pure culture nor genetically manipulated. Here we use a broad-host range bacteriophage to deliver exogenous DNA to an insect endosymbiont and a closely related free-living species. Our results suggest that broad host range bacteriophages can be used to genetically alter insect endosymbionts in their insect host and, as a result, bypass a culturing requirement to genetically alter these bacteria.

## Introduction

Bacteriophages (phages) are the most abundant and diverse biological entities on the planet. With an estimated population size greater than 1×10^31^ (Hendrix *et al*. 1999), these bacterial viruses play essential ecological and evolutionary functions. Phages control the size of bacterial populations and shape the diversity of microbial communities by modulating the abundance of bacterial lineages and promoting, directly and indirectly, the exchange of genetic information among species (Touchon *et al*., 2017; Soucy *et al*. 2015). Historically, phages have played a central role in the development of molecular biology, enabling, for instance, the establishment of DNA as the genetic material of living cells (Hershey and Chase, 1952). Today, phages are widely used as tools in the study of bacteria. For instance, generalized transducing phages such as P1 allow the rapid transfer of DNA among bacterial strains, greatly facilitating genetic dissection of biological processes (Thomason *et al*. 2007).

P1 is a temperate bacteriophage capable of alternating between lytic and lysogenic infection. P1 was initially described in studies involving lysogenic strains of *Escherichia coli* (Bertani, 1951). This phage is capable of mediating generalized transduction (Lennox, 1955), a property that has fostered its adoption as an important experimental tool for the genetic analysis and manipulation of *E. coli* (Thomason *et al*. 2007, Yarmolinsky and Sternberg, 1988). Notably, in addition to its habitual *E. coli* host, P1 can also infect a large number of Gram negative bacterial species (Yarmolinsky and Sternberg, 1988; Goldberg *et al*., 1974; Ornellas and Stocker, 1974; Kaiser and Dworkin, 1975; Murooka and Harada, 1979). This broad host range, along with its well-characterized molecular biology and established experimental procedures, has prompted the use of this phage as an experimental tool for the delivery of DNA to a large number of bacterial species (Streicher *et al*., 1971; O’Connor *et al*., 1983; Downard, 1988; Wolf-Watz *et al*., 1985; Westwater *et al*., 2002; Butela and Lawrence, 2012). Here we establish that P1 is capable of infecting two members of the bacterial genus *Sodalis*, including *Sodalis glossinidius* (Dale and Maudlin, 1999; Chari *et al*., 2015).

*Sodalis glossinidius* is a maternally inherited, Gram negative bacterial endosymbiont of tsetse flies (*Glossina* spp.; Diptera: *Glossinidae*). Similar to other insect endosymbionts, *S. glossinidius* exists in a stable, chronic association with its insect host, and undergoes a predominantly maternal mode of transmission (Aksoy *et al*., 1997; Cheng and Aksoy, 1999; De Vooght *et al*., 2015). Notably, like other insect endosymbionts, this bacterium has undergone an extensive process of genome degeneration as a result of a recent evolutionary transition from free-living existence to permanent host association (McCutcheon *et al*., 2019; Moran *et al*., 2008). Because this process is accompanied by the loss of metabolic capability and stress response pathways (McCutcheon *et al*., 2019; Moran *et al*., 2008; Toh *et al*., 2006; Pontes *et al*. 2011; Clayton *et al*., 2012; Pontes and Dale, 2006), *S. glossinidius* has proven refractory to harsh artificial DNA transformation procedures that are commonly employed in model organisms such as *Escherichia coli* (Pontes and Dale, 2006). Consequently, this bacterium has remained genetically intractable (Kendra *et al.* 2020).

In this study, we demonstrate that the bacteriophage P1 is capable of infecting, lysogenizing and promoting transduction in *Sodalis glossinidius*, and its free-living close relative, the plant-associated and opportunistic pathogen *Sodalis praecaptivus* (Dale and Maudlin, 1999; Chari *et al*., 2015). We demonstrate that P1 can be used to mediate generalized transduction of chromosomal and extrachromosomal DNA in *S. praecaptivus*. We use P1 to transduce autonomous replicating phagemids containing an array of reporter genes and Tn*7* transposition systems harboring fluorescent proteins for chromosomal tagging. Finally, we developed a suicide phagemid containing a mariner transposase for random mutagenesis of bacterial strains susceptible to P1 infection. This study establishes a new efficient method for genetic manipulation of *Sodalis* species (Fig. 1) that can be readily adapted to other Gram negative bacteria. Furthermore, these results provide a potential means for the genetic modification of bacterial endosymbionts, in their insect host, through the use of P1 as a DNA-delivery system.

**Fig. 1.**
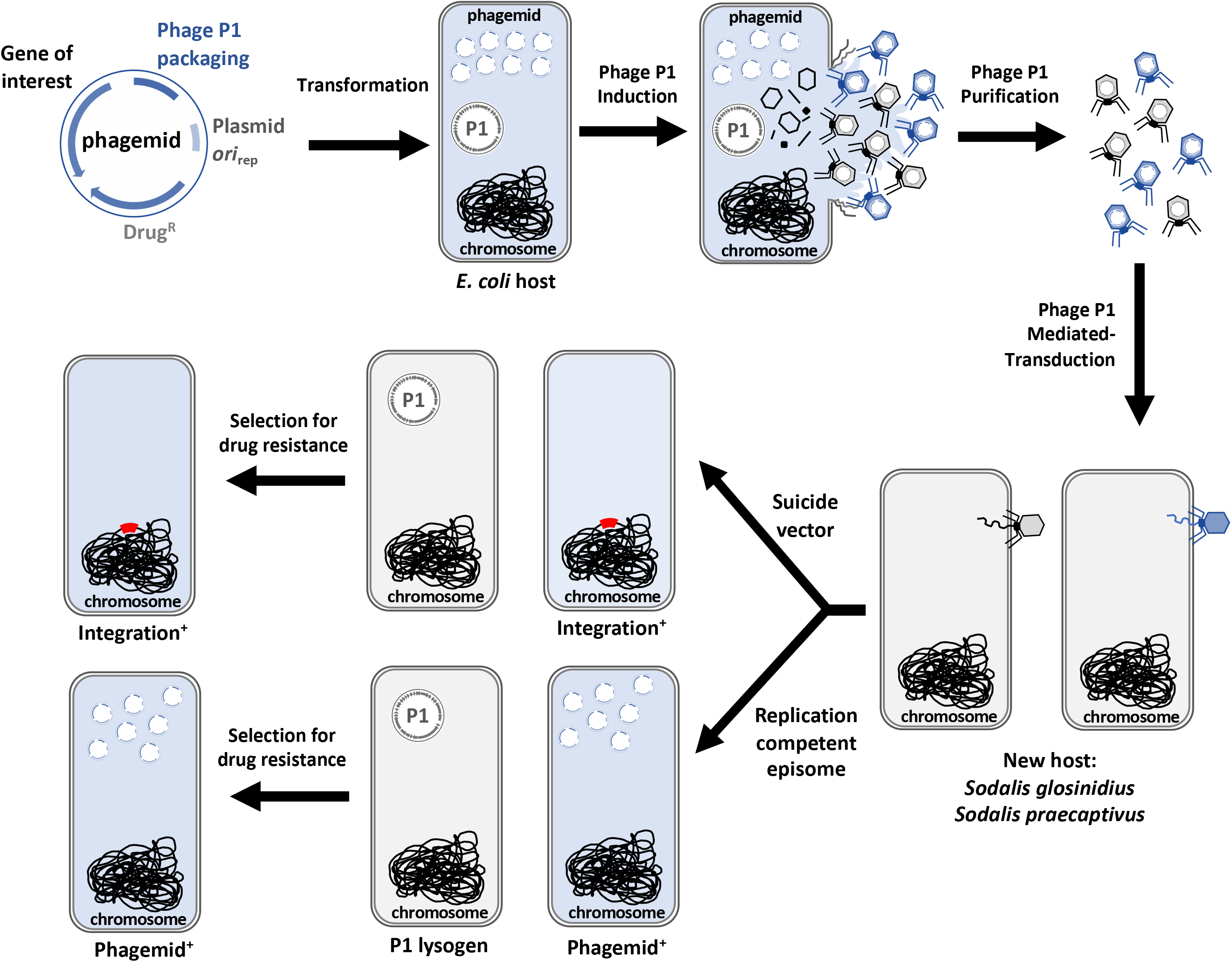
Cartoon representation depicting a workflow of the transduction procedure developed for introduction of phagemids in *Sodalis* species. Following the direction of the arrows: An *E. coli* P1 lysogen host is transformed with a P1 phagemid. The phagemid is packaged following induction of the P1 prophage, and lysates derived from culture supernatant are used to infect a *Sodalis* recipient strain. Cells receiving the phagemid are subsequently isolated on plates containing a selective agent.

## Results

### Bacteriophage P1 infects, lysogenizes and forms phage particles in *Sodalis glossinidius* and *Sodalis praecaptivus*

P1CM*clr*-100(ts) is a thermo-inducible P1 variant harboring a chloramphenicol resistant marker. P1CM*clr*-100(ts) forms chloramphenicol resistant lysogens at low temperatures (≤30°C) but produces phage particles at higher temperatures (≥37°C) (Rosner, 1972). Consequently, infection of *E. coli* by P1CM*clr*-100(ts) yields chloramphenicol resistant lysogens at 30°C. We took advantage of these P1CM*clr*-100(ts) properties to test if *S. glossinidius* and *S. praecaptivus* were susceptible to P1 infection. We exposed cultures of these bacteria to increasing concentrations of P1CM*clr*-100(ts) phage particles and subsequently plated dilutions on solid medium containing chloramphenicol. We established that, similar to the *E. coli* control (Fig. 2A), exposure to increasing concentrations of P1CM*clr*-100(ts) particles yielded increasing numbers of chloramphenicol resistant colonies in both *S. glossinidius* (Fig. 2B) and *S. praecaptivus* (Fig. 2C). Importantly, no chloramphenicol resistant colonies were observed in cultures that were not exposed to P1CM*clr*-100(ts) particles (Fig. 2B and C).

**Fig. 2.**
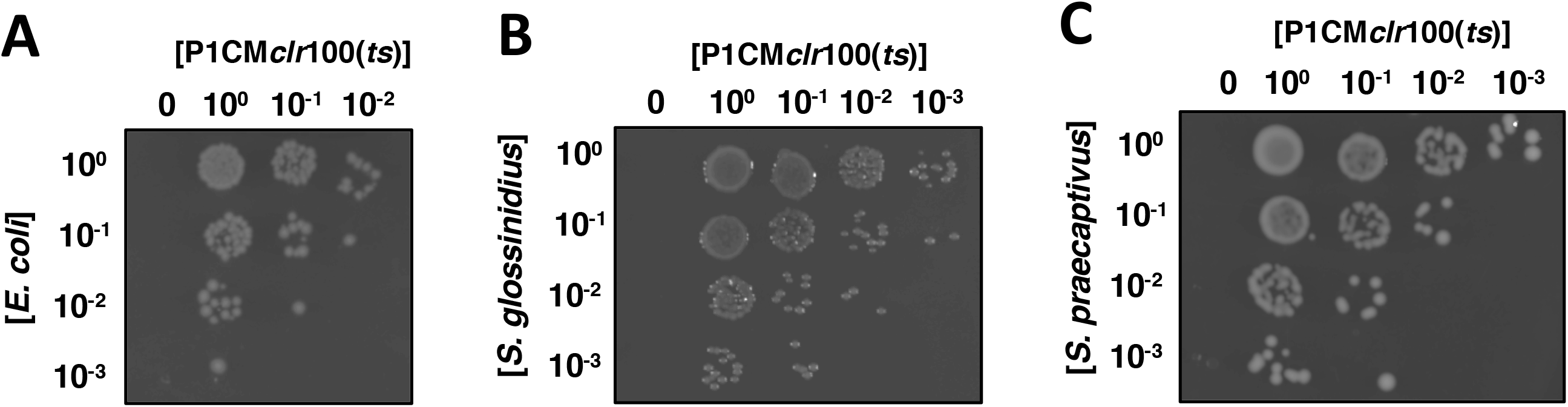
Infection of bacterial strains by phage P1. Lysates derived from an *E. coli* P1CM*clr*-100(ts) lysogen (KL463) were used to infect *E. coli* MG1655 (A), *Sodalis glossinidius* (B) and *Sodalis praecaptivus* (C). Plates depict the formation of chloramphenicol resistant colonies as functions of the concentration of bacteria (vertical axis) and the concentration of P1CM*clr*-100(ts) lysates (horizontal axis). Note that P1 infection conditions for the strains are different (see Materials and Methods), and images do not reflect efficiency of P1 infection. Images show representative plates of at least 3 routine experiments.

That these colonies were P1 lysogens, as opposed to recombinants harboring only the P1-derived chloramphenicol resistant marker, was supported by several lines of evidence. First, the presence of a P1 DNA fragment was detected by polymerase chain reaction (PCR) in both chloramphenicol resistant *S. glossinidius* (Fig. 3A and B) and *S. praecaptivus* clones (Fig. 3C and D), but not in the wild-type strains (Fig. 3A-D, left side). This indicated that chloramphenicol resistant cells harbor at least part of the P1CM*clr*-100(ts) genome. Second, lysates prepared from *S. glossinidius* and *S. praecaptivus* chloramphenicol resistant clones, but not their wild-type counterparts, formed plaques in soft agar cultures of *E. coli* grown at 37°C, a temperature that induces P1CM*clr*-100(ts) lytic replication (Rosner, 1972) (Fig. 3E and F). This indicated that chloramphenicol resistant *S. glossinidius* and *S. praecaptivus* clones can produce phage particles that are lytic to *E. coli* grown at 37°C. Third, the lytic activity of lysates derived from chloramphenicol resistant *S. praecaptivus* cultures propagated at 37°C was 10,000 times higher than those maintained at 30°C (Fig. 3G). This established that higher titers of phage particles were being produced in *S. praecaptivus* chloramphenicol resistant clones at a temperature where P1CM*clr*-100(ts) becomes lytic. Finally, lysates derived from *S. glossinidius* and *S. praecaptivus* chloramphenicol resistant clones, but not their wild-type isogenic counterparts, promoted the formation of chloramphenicol resistant *E. coli* cells at 30°C (Fig. 3H). This indicated that the chloramphenicol resistant marker can be transduced from *S. glossinidius* and *S. praecaptivus* back to *E. coli*.

**Fig. 3.**
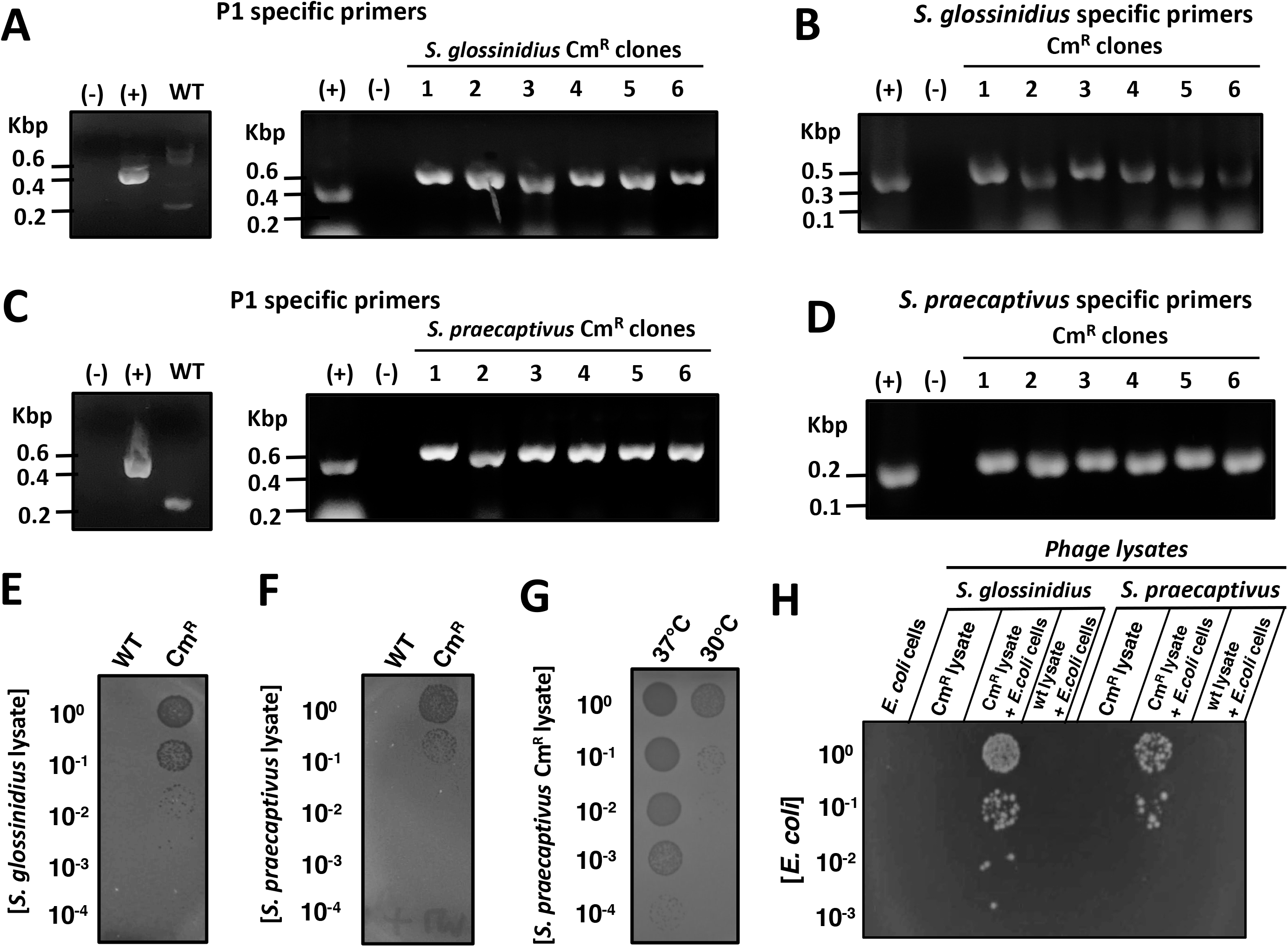
Lysogenization and production of infective phage particles by *Sodalis glossinidius* and *Sodalis praecaptivus* P1 lysogens. (A) Detection of P1 *pacB* gene by PCR and agarose gel electrophoresis in *S. glossinidius* chloramphenicol resistant clones that emerged following exposure to an *E. coli* P1CM*clr*-100(ts) lysogen (KL463). (B) Detection of a *S. glossinidius* specific DNA fragment in clones depicted in (A) by PCR and agarose gel electrophoresis. (C) Detection of P1 *pacB* gene by PCR and agarose gel electrophoresis in *S. praecaptivus* chloramphenicol resistant clones that emerged following exposure to an *E. coli* P1CM*clr*-100(ts) lysogen (KL463). (D) Detection of a *S. praecaptivus* specific DNA fragment in clones depicted in (C) by PCR and agarose gel electrophoresis. (E-G) Formation of phage plaques on soft agar embedded with *E. coli* MG1655. Soft agar plates were spotted with dilutions of lysates derived from wild-type and *S. glossinidius* chloramphenicol resistant (Cm^R^) P1CM*clr*-100(ts) lysogen (MP1705) (E), wild-type and *S. praecaptivus* Cm^R^ P1CM*clr*-100(ts) lysogen (MP1703) (F), and *S. praecaptivus* Cm^R^ P1CM*clr*-100(ts) lysogen (MP1703) grown either at 37℃ or 30℃. Plates are representative of routine experiments. (H) Emergence of Cm^R^ *E. coli* MG1655 following exposure to lysates derived from wild-type *S. glossinidius*, *S. glossinidius* Cm^R^ P1CM*clr*-100(ts) lysogen (MP1705), wild-type *S. praecaptivus*, and *S. praecaptivus* Cm^R^ P1CM*clr*-100(ts) lysogen (MP1703). Plate is representative of at least 3 experiments.

In *S. glossinidius*, the frequency of chloramphenicol resistant colonies arising following P1CM*clr*-100(ts) exposure was similar to those observed for the *E. coli* control cells, indicating that P1 infection occurs efficiently in this bacterium. By contrast, chloramphenicol resistant *S. praecaptivus* colonies emerged at a lower frequency, and higher concentrations of bacterial cells were typically used in P1 infection experiments (see Materials and Methods). Notably, in P1-resistant *Salmonella enterica*, the efficiency of P1 infection can be drastically increased by mutations in either *galU* or *galE* (Ornellas and Stocker, 1974). Because these mutations remove the O-antigen by truncating the core region of the lipopolysaccharide (LPS) (Wilkinson and Stocker, 1968; Osborn, 1968), they presumably facilitate access of P1 to its host receptor—conserved structural motifs within the LPS core (Yarmolinsky and Sternberg, 1988; Ornellas and Stocker, 1974). In particular, while the LPS of *S. praecaptivus* contains structural components attached to its core region, *S. glossinidius* is devoid of such structures (Fig. S1A). Nonetheless, the lower infectivity of P1 does not appear to be related to the physical occlusion of the P1 receptor by components present in the outer portion of the *S. praecaptivus* LPS. This is because a mutation in *galU* results in a truncated LPS in *S. praecaptivus*, but does not affect P1 infectivity (Fig. S1). Hence, unlike *S. enterica*, this phenotype is not due to the presence of P1-antagonizing structure(s) in the outer portion of *S. praecaptivus* LPS. Taken together, these results indicate that phage P1 is capable of infecting and lysogenize in *S. glossinidius* and *S. praecaptivus*.

### P1 generalized transduction in *Sodalis praecaptivus*

During the formation of P1 virions, approximately 0.05-0.5% of infective phage particles package random DNA fragments derived from the bacterial host (Ikeda and Tomizawa, 1965). These particles can mediate the transfer of bacterial DNA across P1 susceptible strains through generalized transduction. In the laboratory, generalized transduction of DNA can be identified by virtue of genetic markers that are packaged in these phage particles and transferred between bacterial strains. Accordingly, we sought to determine if P1 could mediate generalized transduction in *S. praecaptivus*. First, we exposed wild-type *S. praecaptivus* to phage lysates derived from a *S. praecaptivus* P1CM*clr*-100(ts) lysogen harboring the ampicillin resistant (Amp^R^) plasmid pSIM6 (Datta *et al*., 2006). Following lysate exposure, we were able to retrieve Amp^R^ *S. praecaptivus* transductants. Importantly, Amp^R^ cells were absent from both phage lysates alone and cultures of wild-type *S. praecaptivus* that were not exposed to phage (data not shown). In agreement with the notion that these Amp^R^ clones were P1 transductants, diagnostic PCR revealed the presence of a pSIM6 fragment in these cells (Fig. 4A).

**Fig. 4.**
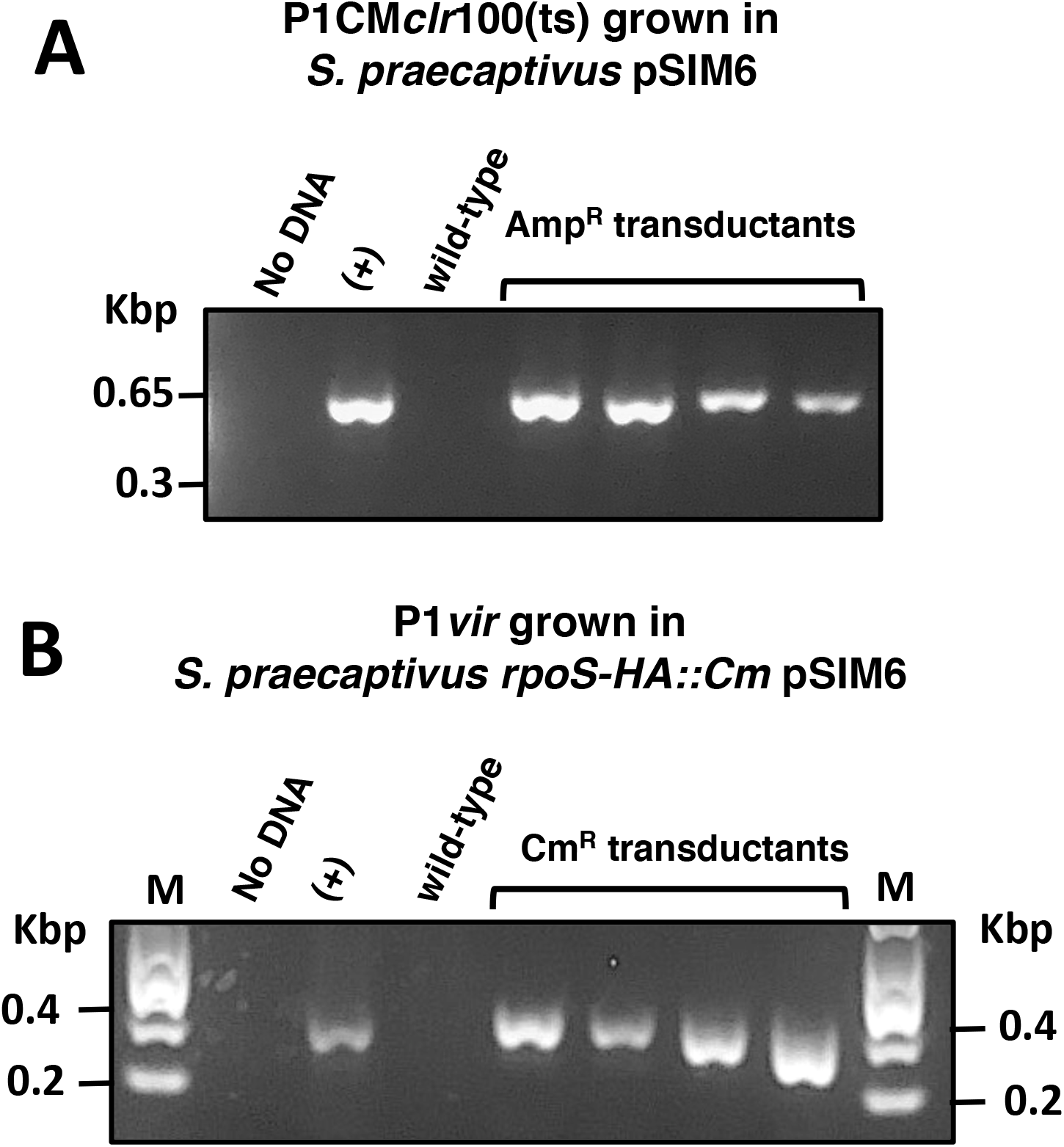
Bacteriophage P1-mediated generalized transduction in *S. praecaptivus*. (A) Detection of a pSIM6 DNA region by PCR and agarose gel electrophoresis in ampicillin resistant (Amp^R^) *S. praecaptivus* transductants following exposure to lysates derived from *S. praecaptivus* Cm^R^ P1CM*clr*-100(ts) (MP1703) harboring Amp^R^ plasmid pSIM6. (B) Detection of the *rpoS*-HA::Cm chromosomal insertion by PCR and agarose gel electrophoresis in *S. praecaptivus* transductants following exposure to lysates of P1*vir* grown in *S. praecaptivus rpoS*-HA::Cm pSIM6 (MP1522) strain.

Next, we attempted to transduce a chromosomal chloramphenicol resistant marker (*rpoS-HA::Cm*) using the P1 lytic strain P1*vir*. [This P1 strain is widely used as a transducing agent in *E. coli* due to its inability to lysogenize cells upon infection and the ease with which transducing lysates can be generated (Thomason *et al*. 2007; Ikeda and Tomizawa, 1965)]. We infected wild-type *S. praecaptivus* cells with P1*vir* lysates grown in a *S. praecaptivus rpoS-HA::Cm* pSIM6 strain. Whereas chloramphenicol resistant (Cm^R^) cells emerged from wild-type *S. praecaptivus* exposed to phage, no Cm^R^ cells were obtained from phage lysates or cultures of naïve wild-type *S. praecaptivus* alone (data not shown). Notably, diagnostic PCR indicated that chloramphenicol resistant clones were transductants harboring the *rpoS-HA::Cm* genetic modification (Fig. 4B). Importantly, all of these *rpoS-HA::Cm* transductants were sensitive to ampicillin, indicating that P1*vir* mediated the transduction of a discrete portion of the *S. praecaptivus* genome. Taken together, these results indicate that bacteriophage P1 can be used to mediate generalized transduction in *S. praecaptivus*.

### Introduction of exogenous DNA in *Sodalis glossinidius* and *Sodalis praecaptivus* by P1-mediated guided transduction

Whereas up to 0.5% of P1 particles can contain random fragments of bacterial host DNA (Ikeda and Tomizawa, 1965), the vast majority of virions harbor P1 DNA. This is because the packaging of P1 genome into phage particles is guided by elements encoded within its DNA sequence (Sternberg and Coulby, 1987a and 1987b). Particularly, this packaging element can be cloned into plasmids (to produce phagemids) or incorporated into the bacterial chromosome to increase the frequency of P1-mediated transduction of adjacent DNA (Westwater *et al*., 2002; Kittleson *et al*., 2012; Huang and Masters, 2014). Indeed, the P1 packaging element can increase the transduction of linked DNA by 1,600-fold above the levels obtained in generalized transduction (Kittleson *et al*., 2012). Hence, this DNA element can be used to increase the number of transducing particles and, consequently, the efficiency of DNA transfer among bacterial strains that are susceptible to P1 infection.

The lack of genetic tools available for the manipulation of *Sodalis* species, specifically *S. glossinidius*, prompted us to explore P1 as a plasmid DNA-delivery tool for these bacteria. As a proof of principle, we used the aforementioned general technique to transfer a number of P1 phagemids (Kittleson *et al*., 2012) (Fig. 5A) into *S. glossinidius*. We were able to recover transductants expressing an array of phenotypic traits encoded in the phagemids. These included light production (*luxCDABE* genes), violacein pigment synthesis (*vioABCE*), β-galactosidase activity (*lacZ*) or green fluorescence (*gfp*) (Fig. 5B and 5C). To expand the tool set available for the modification of *Sodalis* species, we constructed (1) two phagemids for tagging bacterial chromosomes with fluorescent genes at the Tn*7* attachment site (Peters and Craig, 2011), and (2) a suicide phagemid encoding a Himar1 transposition system for random mutagenesis (Rubin *et al*., 1999) (Fig. S2). Following packaging into P1 virions in an *E. coli* P1CM*clr*-100(ts) lysogen, these phagemids were efficiently delivered to *S. glossinidius* and *S. praecaptivus* (Fig. 5D and 5E). Together, these results establish that bacteriophage P1 can be used to efficiently deliver replication-competent and suicide vectors into *S. glossinidius* and *S. praecaptivus* through a “guided transduction” strategy.

**Fig. 5.**
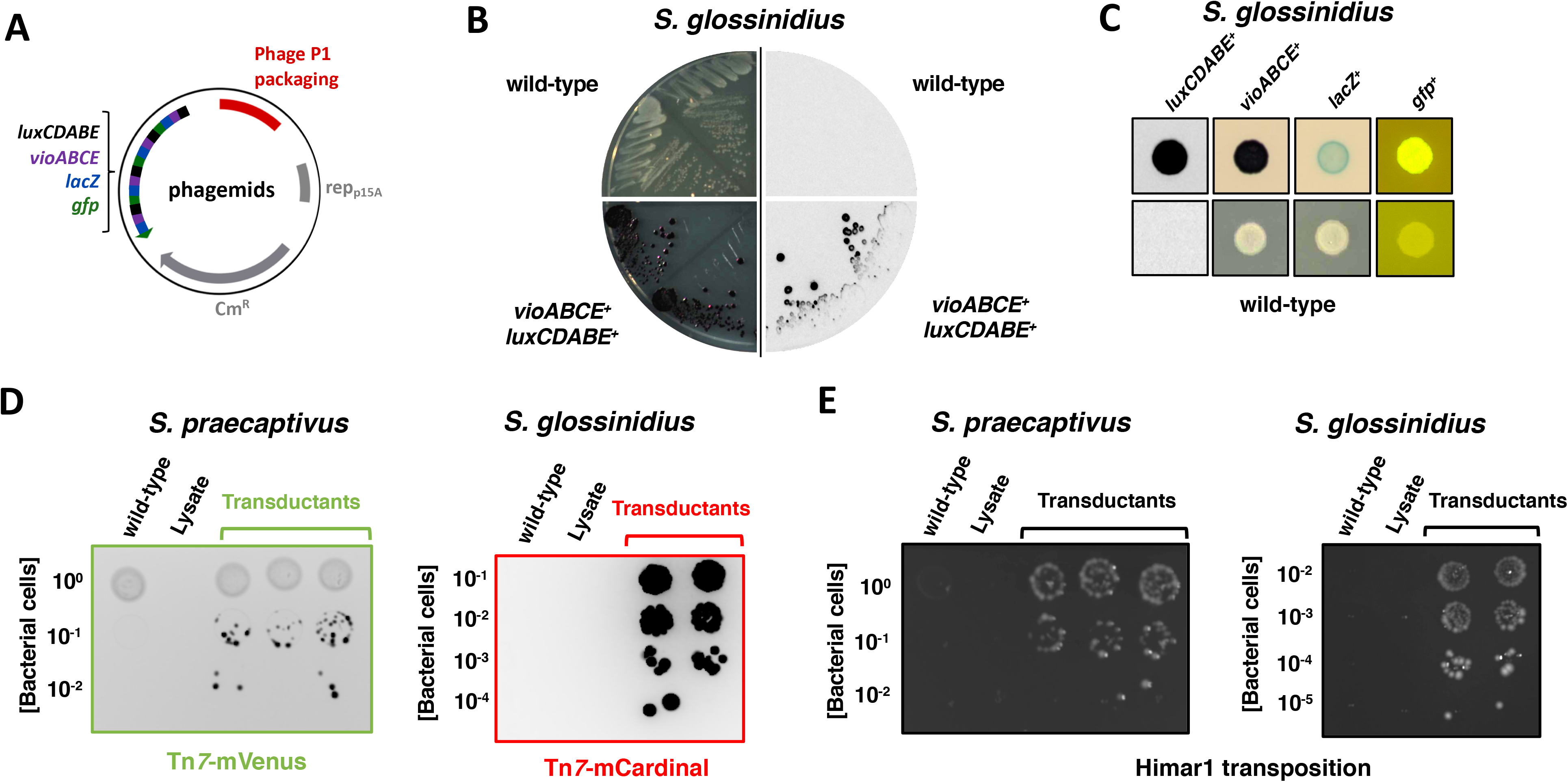
Transduction of P1 phagemids into *Sodalis* species. (A) Schematic representation of replication competent P1 phagemids encoding a number of phenotypic markers (Kittleson *et al*., 2012). (B) Comparison of wild-type (top quadrants) and *S. glossinidius* transductants (lower quadrants) carrying P1 phagemid BBa_J72114-BBa_J72104 (Kittleson *et al*., 2012). Transductant colonies are purple due to the expression of violacein biosynthetic genes (lower, left quadrant), and produce light due to the expression of bioluminescence genes (lower, right quadrant). (C) Macrocolonies derived from wild-type *S. glossinidius* (bottom row) and transductants carrying P1 phagemids BBa_J72114-BBa_J72104 (*luxCDABE*^+^ and *vioABCE^+^*), BBa_J72114-BBa_J72100 (*lacZ^+^*) or BBa_J72113-BBa_J72152 (*gfp^+^*) (upper row). (D) Transduction of pP1-Tn*7*-mVenus into *S. praecaptivus* (left-hand side plate) and pP1-Tn*7*-mCardinal into *S. glossinidius* (right-hand side plate). (E). Transduction of phagemid encoding a Himar1 transposition system (pP1-Himar) into *S. praecaptivus* (left-hand side plate) and *S. glossinidius* (right-hand side plate).

## Discussion

In the current study, we demonstrate that bacteriophage P1 can infect, lysogenize and promote transduction in two species of the genus *Sodalis*. We show that P1 can mediate generalized transduction in *S. praecaptivus* (Fig. 4), and we establish that this bacteriophage can be used for the delivery of plasmids and suicide vectors for the genetic manipulation of *S. glossinidius* and *S. praecaptivus* (Fig. 5). While these results constitute a significant advance in the development of genetic modification tools to study these bacterial species, they also clear the way for the implementation of P1-based DNA-delivery systems to uncultured *Sodalis* species (Heddi *et al*., 1998; Fukatsu *et al*., 2007; Nováková and Hypsa, 2007; Chrudimský *et al*., 2012; Smith *et al*., 2013; Santos-Garcia *et a*l., 2017; Šochová *et al*., 2017) and Gram negative insect endosymbionts belonging to other genera.

Whereas *S. praecaptivus* can be genetically engineered with relative ease, the ability of P1 to mediate generalized transduction provides a number of applications for the manipulation of this bacterium. For instance, although *S. praecaptivus* can be readily modified by recombineering functions of phage λ (λ-Red; Clayton *et al.* 2017; Enomoto *et al*., 2017; Thomason *et al*., 2014), the use of this technique has two major drawbacks. First, the expression of recombineering functions can be mutagenic (Murphy and Campellone, 2003). This can potentially produce confounding results in subsequent experiments as phenotypes associated with a particular engineered modification may actually results from secondary mutation(s). Second, typical temperature-sensitive plasmids (rep_pSC101_^ts^ *ori*) harboring recombineering functions cannot be cured from *S. praecaptivus* by propagating cells at non-permissive temperatures (≥37°C) in the absence of plasmid selection (our unpublished results). The inability to cure these plasmids can increase the chances of secondary mutations through leaky expression of recombineering functions, and hinder the use of plasmids from the same incompatibility group in downstream genetic analyses. Importantly, both of these issues can be overcome by P1-mediated generalized transduction. That is, genomic DNA fragments engineered using λ-Red can be transferred to naïve *S. praecaptivus* cells that lack recombineering plasmids and, therefore, have not been exposed to potential mutagenic events (Fig. 4B).

By contrast, the establishment of P1 mediated transduction provides a considerable advancement in our ability to genetically manipulate *S. glossinidius*. This is because *S. glossinidius* is recalcitrant to DNA transformation by standard techniques such as heat-shock and electroporation (Pontes and Dale, 2006 and 2011; Kendra *et al*., 2020). Whereas we have recently developed a method for DNA transfer to *S. glossinidius* via conjugation (Kendra *et al*., 2020), P1-mediated transduction provides an alternative, simpler method for the introduction of exogenous DNA into this bacterium. Altered chromosomal fragments, replication competent plasmids carrying an array of functions, and suicide vectors engineered for allelic replacement or containing transposition systems can be quickly transduced into *S. glossinidius* in a simple protocol. Given the large DNA packaging capability of P1 (up to 100 Kbp) (Thomason *et al*. 2007), this bacteriophage can be efficiently used for a variety of applications, including the delivery of bacterial artificial chromosomes (BAC vectors) or large plasmids encoding multiple genome editing CRISPR systems (Adiego-Pérez *et al*., 2019) that are not easily transferred by conjugation (Kendra *et al*., 2020). Additionally, the P1 packing sequence can be incorporated into DNA fragments used in insertional mutagenesis (Huang and Masters, 2014), enabling rapid and efficient combination of mutations via P1 “guided transduction.” This approach can greatly facilitate the implementation of several analyses (e.g. complementation and epistasis) to identify and dissect genetic components and pathways governing bacterial behaviors and interactions with eukaryotic hosts.

Beyond the genus *Sodalis*, the results highlighted in this study have potential broad implications for the genetic modification of uncultured Gram negative insect endosymbionts. In Gram negative bacteria, the LPS is the major structural constituent of the outer leaflet of the outer membrane. The LPS is composed of a highly conserved lipid A “anchor,” a conserved core polysaccharide and, sometimes, a hypervariable outer component designated O-antigen (Fig. S1A; Silipo and Molinaro, 2010). Bacteriophage P1 has a broad host range, in part, because it recognizes, as its host receptor, structural features of the conserved LPS core (Yarmolinsky and Sternberg, 1988). In addition to *E. coli* and several species of the Gammaproteobacteria, P1 has been shown to be capable of infecting various members within the Alpha, Beta- and Deltaproteobacteria, and even bacterial species residing outside the proteobacteria Phylum such as *Flavobacterium* sp. M64 (Yarmolinsky and Sternberg, 1988; Goldberg *et al*., 1974; Ornellas and Stocker, 1974; Kaiser and Dworkin, 1975; Murooka and Harada, 1979).

Notably, a large number of economically important insect species—including several disease vectors of animals and plants—harbor maternally inherited, Gram negative bacterial endosymbionts (McCutcheon *et al*., 2019; Moran *et al*., 2008). However, unlike *S. glossinidius* and a handful of other species, the vast majority of these endosymbionts have not been isolated in pure culture (Dale and Maudlin, 1999; Chrudimský *et al*., 2012; Hypsa and Dale, 1997; Sabri *et al*., 2011; Dale *et al*., 2006; Brandt *et al*., 2017; Masson *et al*., 2018). This is because these bacteria undergo a process of genome degeneration and size reduction during the course of long-term evolution and specialization within their eukaryotic insect hosts. This process leads to the loss of many physiological functions that are required for replication outside the host (McCutcheon *et al*., 2019; Moran *et al*., 2008; Pontes *et al*., 2011; Pontes and Dale, 2006). Notably, classical protocols of bacterial genetics require the manipulation of large numbers of cells and the subsequent isolation of rare genetic events as bacterial colonies on selective agar plates. Consequently, the implementation of genetics for the study of insect endosymbionts has remained scarce and limited to species that can grow in axenic culture, form colonies on agar plates and are receptive to exogenous DNA (Pontes and Dale, 2006; Kendra *et al*., 2020; Masson *et al*., 2018).

The ability of bacteriophage P1 to deliver DNA to a large number of bacterial species, suggests a clear method in which the requirement for culturing may be bypassed. That is, similar to viral vectors that are commonly utilized in gene therapy in mammalians (Kay *et al*., 2011), P1 could be used to deliver DNA to bacterial endosymbionts inside their insect hosts. Specifically, insects could be microinjected (De Vooght *et al*., 2015; Pontes *et al*., 2011; Boyle *et al*., 1993; Oliver *et al*., 2003; Doremus *et al*., 2018) with P1 virions packaged with recombinant DNA. The establishment of successful P1 infections and subsequent enrichment of transductant endosymbionts to near-homogeneous or clonal populations could be attained by making use of phenotypic markers (Fig. 5) and implementing antibiotic selection regiments in insects (Wilkinson, 1998; Dale and Welburn, 2001; Dobson and Rattanadechakul, 2001; Koga *et al*., 2007; Pais *et al*., 2008; McLean *et al*., 2011). Whereas this approach would preferentially target recently acquired endosymbionts by virtue of their ability to exist intra and extracellularly in various insect tissues (Aksoy *et al*., 1997; Cheng and Aksoy, 1999; De Vooght *et al*., 2015; Moran *et al*., 2008), ancient obligate intracellular endosymbionts could also be subjected to infection and genetic modification via P1, if they transiently exit host cells (Zaidman-Rémy *et al*., 2018).

The results presented in this study pave the way for the development of tractable genetic systems for *S. glossinidius* and, potentially, a myriad of Gram negative bacterial endosymbionts of insects. While this may empower the use of genetics to study these obscure bacteria, it has also clear translational applications. P1-mediated DNA delivery into insect endosymbionts may allow the engineering of bacterial traits aimed at modifying aspects of insect ecology (Pontes and Dale, 2006; Kendra *et al*., 2020), mitigating their burden on economic activities and human health.

## Material and Methods

### Microbial strains, phages, plasmids and growth conditions

Microbial strains, phages and plasmids used in this study are presented in Table S1. Unless indicated, all *E. coli* strains were propagated at 30, 37 or 42°C in Luria Bertani (LB) broth or agar (1.5% w/v). *Sodalis glossinidius* was grown at 27°C in brain-heart infusion broth supplemented with 10 mM MgCl_2_ (BHI) or on brain-heart infusion agar (1.2% w/v) supplemented with 10 mM MgCl_2_ (BHI agar). *Sodalis glossinidius* was also propagated on BHI agar plates supplemented with 10% defibrinated horse blood (BHIB). *Sodalis praecaptivus* was grown at 30, 39 or 42°C in Luria Bertani (LB) broth or agar (1.5% w/v) lacking sodium chloride. For experiments involving P1 infection or generation of lysates, the growth media was supplemented with CaCl_2_ and MgCl_2_ to a final concentration of 10 mM respectively. Growth of *Sodalis glossinidius* on BHIB agar plates was carried out under microaerophilic conditions, which was achieved either using BD GasPak EZ Campy Gas Generating sachets or a gas mixture (5% oxygen 95% CO_2_). For all strains, growth in liquid medium was carried out in shaking water bath incubators with aeration (250 rpm). When required, medium was supplemented with ampicillin (100 μg/mL), chloramphenicol (20 μg/mL for *E. coli* or *S. praecaptivus*, and 10 μg/mL for *S. glossinidius*), kanamycin (50 μg/mL for *E. coli* and 25 μg/mL for *S. glossinidius* or *S. praecaptivus*). Arabinose was used at a concentration of 0.5 or 1% (w/v); 5-bromo-4-chloro-3-indolyl-β-D-galactopyranoside (X-gal) was used at a concentration of 100 μg/mL.

### Lipopolysaccharide extraction and detection

Extraction of lipopolysaccharide (LPS) from *S. glossinidius* and *S. praecaptivus* cultures were carried out as described (Davis and Goldberg, 2012). Extracted samples were separated in a NuPAGE 10% Bis-Tris gel in NuPAGE MES SDS Running (ThermoFisher Scientific). LPS in gels were stained with ProteoSilver Silver Stain Kit (Sigma Aldrich).

### Construction of phagemid pP1-Tn *7*-mCardinal

Oligonucleotide sequences used in this study are presented in Table S2. Phusion® High-Fidelity DNA Polymerase (New England BioLabs) was used in PCR reactions with primers 469 and 470 and plasmid BBa_J72113-BBa_J72152 (Kittleson *et al*., 2012) as template. The PCR product was ligated into pMRE-Tn*7*-163 (Schlechter *et al*., 2018), previously digested with SbfI, using NEBuilder® HiFi DNA Assembly (New England BioLabs). The integrity of the construct was verified by DNA sequencing and the ability to be efficiently transduced by P1 particles.

### Construction of phagemid pP1-Tn *7*-mVenus

Oligonucleotide sequences used in this study are presented in Table S2. Phusion® High-Fidelity DNA Polymerase (New England BioLabs) was used in PCR reactions with primers 469 and 470 and plasmid BBa_J72113-BBa_J72152 (Kittleson *et al*., 2012) as template. The PCR product was ligated into pMRE-Tn*7*-166 (Schlechter *et al*., 2018), previously digested with SbfI, using NEBuilder® HiFi DNA Assembly (New England BioLabs). The integrity of the construct was verified by DNA sequencing and the ability to be efficiently transduced by P1 particles.

### Construction of phagemid pP1-Himar

Oligonucleotide sequences used in this study are presented in Table S2. Phusion® High-Fidelity DNA Polymerase (New England BioLabs) was used in PCR reactions with primers 475 and 476 and plasmid BBa_J72113-BBa_J72152 (Kittleson *et al*., 2012) as template. The PCR product was ligated into pMarC9-R6k (Lee *et al*., 2019), previously digested with EcoRI and HinDIII, using NEBuilder® HiFi DNA Assembly (New England BioLabs). The integrity of the construct was verified by DNA sequencing and the ability to be efficiently transduced by P1 particles.

### Recombineering procedure for *Sodalis praecaptivus*

Oligonucleotide sequences used in this study are presented in Table S2. A *S. praecaptivus* strain harboring plasmid pSIM6 (Datta *et al*., 2006) was grown overnight in LB broth supplemented with 100 μg/ml of ampicillin at 30°C and 250 rpm. Cells were diluted (1:100) in 30 ml of the same medium and grown to an OD_600_ between 0.45-0.5. The culture flask was then grown in a water bath at 42°C and 250 rpm for 25 min. Cells were immediately transferred to a 50 ml conical tube, collected by centrifugation (7,000 rpm for 2.5 min at 4°C), and resuspended in 40 ml of ice-cold dH_2_O. Cells were collected again by centrifugation, and this washing procedure was repeated a second time. Finally, cells were resuspended in 150 μl of ice-cold dH_2_O. Homologous recombination was obtained by electroporating 70 μl of cell suspension with 10 μl of purified PCR products generated with primers 251 and 252 (*galU::Kn*) or 84 and 85 (*rpoS-HA::Cm*) and plasmids pKD4 and pKD3 (Datsenko and Wanner, 2000) as templates, respectively.

### Preparation of phage lysates derived from EMG16 P1 lysogens

Lysates from *E. coli* EMG16 harboring selected phagemids were prepared following arabinose induction as described (Kittleson *et al*., 2012).

### Preparation of stocks P1 *vir* phage lysates

Stocks of P1*vir* phage lysates were prepared by infecting *E. coli* MG1655 (Blattner *et al*., 1997), as described (Thomason *et al*. 2007).

### Preparation of phage lysates derived from P1CM *clr*-100(ts) lysogens

*Escherichia coli* P1CM*clr*-100(ts) lysogens were grown overnight at 30°C. Cultures were diluted (1:100, v/v) into fresh medium and grown to and OD_600_ value of 0.3-0.4. Subsequently, cultures were shifted to 42°C and propagated until extensive cell lysis (3-4 h). At times, cultures were allowed to grow at 42°C for 16 h prior to the preparation of lysates. Partially lysed cells were disrupted by vortexing the cultures following the addition of chloroform (1 volume of chloroform per 100 volumes of culture). Cell debris was removed by centrifugation (12 min, 4,000 x rpms, room temperature), and the supernatant was passed through a 0.22 μm polyethersulfone membrane filter. *Sodalis praecaptivus* P1CM*clr*-100(ts) lysogens were prepared as described above, except that cells were grown for 2 h at 42°C and 16 h at 37°C prior to lysate preparation. *Sodalis glossinidius* P1CM*clr*-100(ts) lysogens were grown in BHI to an OD600 of 0.4. Cultures were heat shocked at 37 °C for 2 h. The cultures were treated with chloroform (1 volume of chloroform per 100 volumes of culture) and processed as described for *E. coli* and *S. praecaptivus*.

### Infection by P1 *vir*, P1CM *clr*-100(ts) and P1 transducing particles

Following overnight grow in LB, *E. coli* cultures were diluted in fresh LB supplemented with 10 mM CaCl_2_ and MgCl_2_ to an OD_600_ of 1. 1 mL aliquots of these cell solutions were incubated for 30 minutes at 30°C in the absence or presence of various concentrations of P1 lysate. Cells were subsequently collected by centrifugation (1 min, 13,000 x rpms, room temperature), and the supernatants were replaced by 1 ml of LB containing 5 mM sodium citrate. Cells were grown for 1 h at 30°C and 250 rpm prior to plating. *Sodalis praecaptivus* cells grown overnight in LB were collected by centrifugation (1 min, 13,000 x rpms, room temperature) and resuspended in fresh LB supplemented with 10 mM CaCl_2_ and MgCl_2_. One ml of these resuspended solutions was incubated for 30 minutes at 30°C in the absence or presence of various concentrations of P1 lysate. Cells were collected by centrifugation (1 min, 13,000 x rpms, room temperature), and the supernatants were replaced by 1 ml of LB containing 5 mM sodium citrate. Cells were plated following 1 h of growth at 30°C and 250 rpm. *Sodalis glossinidius* cells were grown in BHI for 3-5 days, to an OD_600_ ≈ 0.5. Cells were collected by centrifugation and concentrated to an OD_600_ of 1. One ml of concentrated cultures was incubated for 60 minutes at 30°C in the absence or presence of various concentrations of P1 lysate. Cells were collected by centrifugation (1 min, 13,000 x rpms, room temperature), and the supernatants were replaced by 10 ml of BHI. Cultures were incubated overnight at 27°C overnight with shaking prior to plating.

### Curing of pP1-Tn *7* phagemids

Transduction of pP1-Tn*7* phagemids into *S. glossinidius* and *S. praecaptivus* were initially selected on plates containing ampicillin (Fig. 5D, Fig. S2). Because episomes harboring rep_pSC101_^ts^ origins of replication are not easily cured from *S. praecaptivus* (see discussion), the curing of phagemids was only performed in *S. glossinidius*. The strategy used to identify *S. glossinidius* clones lacking phagemids was similar to the one adopted elsewhere (Pontes and Dale, 2001). Briefly, to identify *S. glossinidius* clones that contained the chloramphenicol resistant marker at the Tn*7* attachment site and had loss the ampicillin resistant plasmid, cultures were propagated in BHI containing chloramphenicol and 1% arabinose. After 4 passages, cells were diluted and plated. Single colonies were screened for sensitivity to ampicillin. Transposon insertion at the Tn*7* attachment site was verified by PCR with primers 1018 and 1019.

### Image Acquisition, Analysis and Manipulation

DNA agarose gel electrophoresis and bacterial colonies, with the exception of *S. glossinidsius* macrocolonies (were detected using an Amersham Imager 680 (GE Healthcare). *Sodalis glossinidsius* macrocolonies expressing GFP were detected using a dark reader (Clare Chemical Research) and documented with an iPhone. When oversaturated, the intensity of signals in images were adjusted across the entire images using Preview (Apple).

## Acknowledgement

We would like to thank Serap Aksoy (Yale University) for kindly providing us with a culture of *Sodalis glossinidius*, and Hubert Salvail (Yale University) for assistance obtaining an *E. coli* P1CM*clr*-100(ts) lysogen. MHP is supported by grant AI148774 from the National Institutes of Health and startup funds from The Pennsylvania State University College of Medicine.

## Supplemental Material

Figure S1.

Figure S2.

Table S1.

Table S2.

## Figure Legends

**Fig. S1.**
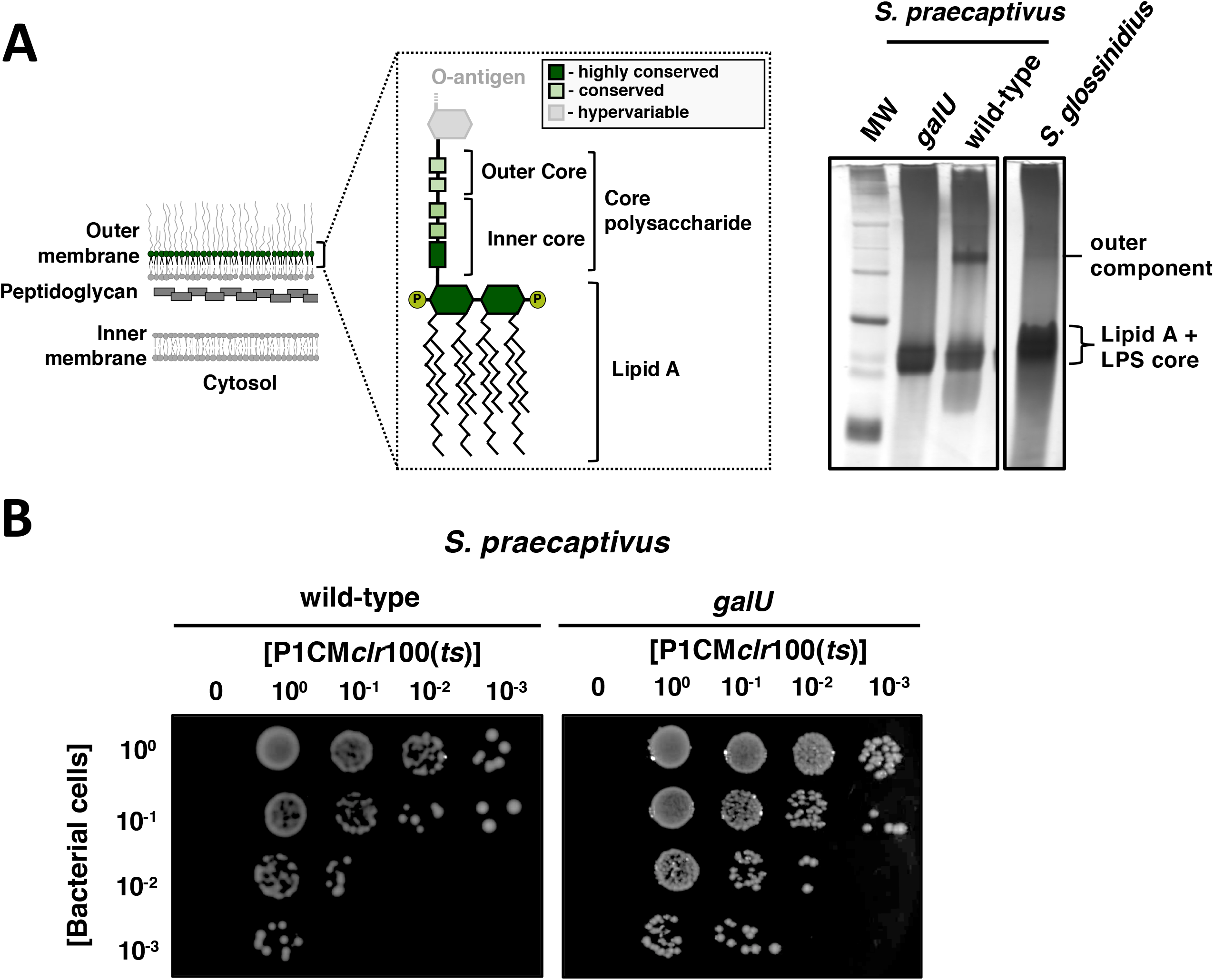
Effect of a *galU* mutation on LPS composition and P1 infectivity in *S. praecaptivus*. (A) Schematic representation depicting the inner membrane, the peptidoglycan and the outer membrane of a prototypical Gram negative bacterium. Inlet shows a more detailed schematic of structural components of the LPS (left-hand side). Silver-stained SDS-PAGE of LPS purified from *S. praecaptivus galU* (CMK36), wild-type *S. praecaptivus* and wild-type *S. glossinidius* (right-hand side). (B) Plates depicting the appearance of chloramphenicol resistant colonies as functions of the concentration of bacteria (vertical axis) and the concentration of P1CM*clr*-100(ts) lysates (horizontal axis). Cm^R^ colonies emerge at similar frequencies in wild-type *S. praecaptivus* (left-hand side plate) and *S. praecaptivus galU* (CMK36) (right-hand side plate) when cells are exposed to equal amounts of P1CM*clr*-100(ts) particles.

**Fig. S2.**
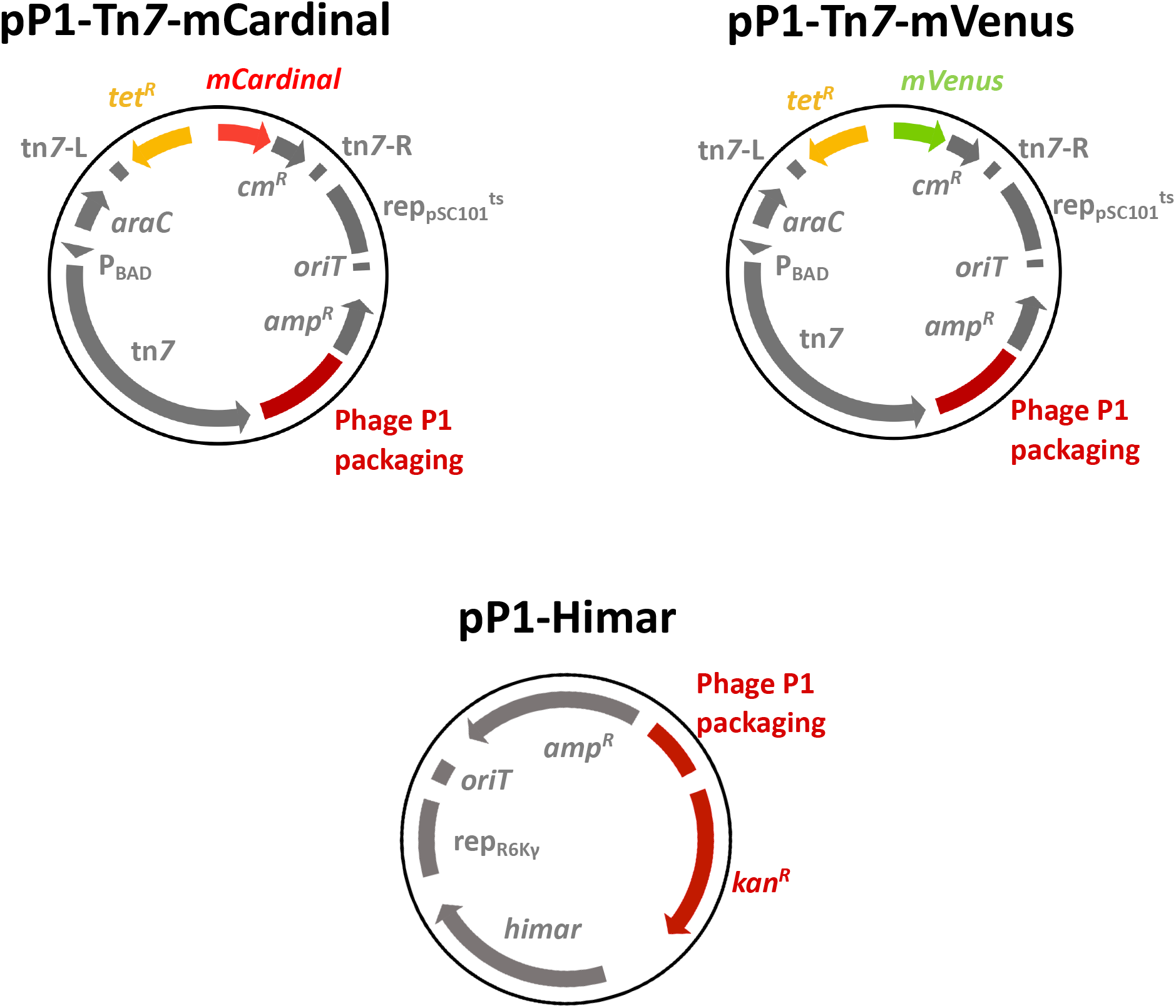
Schematic representation of P1 phagemids used for tagging the bacterial chromosome at the Tn*7* attachment site(s) with genes encoding the fluorescent proteins mCardinal (pP1-Tn*7*-mCardinal) or mVenus (pP1-Tn*7*-mVenus); and the suicide P1 phagemid for random mutagenesis with a Himar1 transposition system (pP1-Himar).

## References

Adiego-Pérez, B., Randazzo, P., Daran, J.M., Verwaal, R., Roubos, J.A., Daran-Lapujade, P., and van der Oost, J. (2019) Multiplex genome editing of microorganisms using CRISPR-Cas. FEMS Microbiol Lett 366:fnz086.

Aksoy, S., Chen, X., and Hypsa, V. (1997) Phylogeny and potential transmission routes of midgut-associated endosymbionts of tsetse (Diptera:Glossinidae). Insect Mol Biol 6:183–190.

Bertani, G. (1951) Studies on lysogenesis. I. The mode of phage liberation by lysogenic *Escherichia coli*. J Bacteriol 62:293–300.

Blattner, F.R., Plunkett, G. 3rd, Bloch, C.A., Perna, N.T., Burland, V., Riley, M., et al. (1997) The complete genome sequence of *Escherichia coli* K-12. Science 277:1453–1462.

Boyle, L., O’Neill, S.L., Robertson, H.M., and Karr, T.L. (1993) Interspecific and intraspecific horizontal transfer of *Wolbachia* in *Drosophila*. Science 260:1796–1799.

Brandt, J.W., Chevignon, G., Oliver, K.M., and Strand, M.R. (2017) Culture of an aphid heritable symbiont demonstrates its direct role in defence against parasitoids. Proc Biol Sci 284:20171925.

Butela, K., and Lawrence, J.G. (2012) Genetic manipulation of pathogenicity loci in non-Typhimurium *Salmonella*. J Microbiol Methods 91:477–482.

Chari, A., Oakeson, K.F., Enomoto, S., Jackson, D.G., Fisher, M.A., and Dale, C. (2015) Phenotypic characterization of *Sodalis praecaptivus* sp. nov., a close non-insect-associated member of the *Sodalis*-allied lineage of insect endosymbionts. Int J Syst Evol Microbiol 65:1400–1405.

Cheng, Q., and Aksoy, S. (1999) Tissue tropism, transmission and expression of foreign genes *in vivo* in midgut symbionts of tsetse flies. Insect Mol Biol 8:125–132.

Chrudimský, T., Husník, F., Nováková, E., and Hypša, V. (2012) *Candidatus* Sodalis melophagi sp. nov.: phylogenetically independent comparative model to the tsetse fly symbiont *Sodalis glossinidius*. PLoS One 7:e40354.

Clayton, A.L., Oakeson, K.F., Gutin, M., Pontes, A., Dunn, D.M., von Niederhausern, A.C., et al. (2012) A novel human-infection-derived bacterium provides insights into the evolutionary origins of mutualistic insect-bacterial symbioses. PLoS Genet 8:e1002990.

Clayton, A.L., Enomoto, S., Su, Y., and Dale, C. (2017) The regulation of antimicrobial peptide resistance in the transition to insect symbiosis. Mol Microbiol 103:958–972.

Dale, C., and Maudlin, I. (1999) *Sodalis* gen. nov. and *Sodalis glossinidius* sp. nov., a microaerophilic secondary endosymbiont of the tsetse fly *Glossina morsitans morsitans*. Int J Syst Bacteriol 49:267–275.

Dale, C., and Welburn, S.C. (2001) The endosymbionts of tsetse flies: manipulating host-parasite interactions. Int J Parasitol 31:628–631.

Dale, C., Beeton, M., Harbison, C., Jones, T., and Pontes M. (2006) Isolation, pure culture, and characterization of “*Candidatus* Arsenophonus arthropodicus,” an intracellular secondary endosymbiont from the hippoboscid louse fly *Pseudolynchia canariensis*. Appl Environ Microbiol 72:2997–3004.

Datsenko, K.A., Wanner, B.L. (2000) One-step inactivation of chromosomal genes in *Escherichia coli* K-12 using PCR products. Proc Natl Acad Sci USA 97:6640–6645.

Datta, S., Costantino, N., and Court, D.L. (2006) A set of recombineering plasmids for gram-negative bacteria. Gene 379:109–115.

Davis, M.R. Jr., and Goldberg, J.B. (2012) Purification and visualization of lipopolysaccharide from Gram-negative bacteria by hot aqueous-phenol extraction. J Vis Exp 63:3916.

De Vooght, L., Caljon, G., Van Hees, J., Van Den Abbeele, J. (2015) Paternal transmission of a secondary symbiont during mating in the viviparous tsetse fly. Mol Biol Evol 32:1977–1980.

Dobson, S.L., and Rattanadechakul, W. (2001) A novel technique for removing *Wolbachia* infections from *Aedes albopictus* (Diptera: Culicidae). J Med Entomol 38:844–849.

Doremus, M.R., Smith, A.H., Kim, K.L., Holder, A.J., Russell, J.A., and Oliver, K.M. (2018) Breakdown of a defensive symbiosis, but not endogenous defences, at elevated temperatures. Mol Ecol 27:2138–2151.

Downard, J.S. (1988) Tn*5*-mediated transposition of plasmid DNA after transduction to *Myxococcus xanthus*. J Bacteriol 170:4939–4941.

Enomoto, S., Chari, A., Clayton, A.L., and Dale, C. (2017) Quorum sensing attenuates virulence in *Sodalis praecaptivus*. Cell Host Microbe 21:629–636.e5.

Fukatsu, T., Koga, R., Smith, W.A., Tanaka, K., Nikoh, N., Sasaki-Fukatsu, K., Yoshizawa, K., et al. (2007) Bacterial endosymbiont of the slender pigeon louse, *Columbicola columbae*, allied to endosymbionts of grain weevils and tsetse flies. Appl Environ Microbiol 73:6660–6668.

Goldberg, R.B., Bender, R.A., Streicher, S.L. (1974) Direct selection for P1-sensitive mutants of enteric bacteria. J. Bacteriol. 118:810–814.

Heddi, A., Charles, H., Khatchadourian, C., Bonnot, G., and Nardon, P. (1998) Molecular characterization of the principal symbiotic bacteria of the weevil *Sitophilus oryzae*: a peculiar G+C content of an endocytobiotic DNA. J Mol Evol 47:52–61.

Hendrix, R.W., Smith, M.C., Burns, R.N., Ford, M.E., Hatfull, G.F. (1999) Evolutionary relationships among diverse bacteriophages and prophages: all the world’s a phage. Proc Natl Acad Sci USA 96:2192–2197.

Hershey, A.D., and Chase, M. (1952) Independent functions of viral protein and nucleic acid in growth of bacteriophage. J Gen Physiol 36:39–56.

Huang, H., and Masters, M. (2014) Bacteriophage P1 *pac* sites inserted into the chromosome greatly increase packaging and transduction of *Escherichia coli* genomic DNA. Virology 468-470:274–282.

Hypsa, V., and Dale, C. (1997) *In vitro* culture and phylogenetic analysis of “*Candidatus* Arsenophonus triatominarum,” an intracellular bacterium from the triatomine bug, *Triatoma infestans*. Int J Syst Bacteriol 47:1140–1144.

Ikeda, H., and Tomizawa, J.I. (1965) Transducing fragments in generalized transduction by phage P1. I. Molecular origin of the fragments. J Mol Biol 14:85–109.

Kaiser, D., and Dworkin, M. (1975) Gene transfer to myxobacterium by *Escherichia coli* phage P1. Science 187:653–653.

Kay, M.A., Glorioso, J.C., and Naldini, L. (2011) Viral vectors for gene therapy: the art of turning infectious agents into vehicles of therapeutics. Nat Med 7:33–40.

Kendra, C.G., Keller, C.M., Bruna, R.E., and Pontes, M.H. (2020) Conjugal DNA transfer in the maternally inherited symbiont of tsetse flies *Sodalis glossinidius*. mSphere 5:e00864–20.

Kittleson, J.T., DeLoache, W., Cheng, H.Y., and Anderson, J.C. (2012) Scalable plasmid transfer using engineered P1-based phagemids. ACS Synth Biol 1:583–589.

Koga, R., Tsuchida, T., Sakurai, M., and Fukatsu, T. (2007) Selective elimination of aphid endosymbionts: effects of antibiotic dose and host genotype, and fitness consequences. FEMS Microbiol Ecol 60:229–239.

Lee, H.H., Ostrov, N., Wong, B.G., Gold, M.A., Khalil, A.S., Church, G.M. (2019) Functional genomics of the rapidly replicating bacterium *Vibrio natriegens* by CRISPRi. Nat Microbiol 4:1105–1113.

Lennox, E.S. (1955) Transduction of linked genetic characters of the host by bacteriophage P1. Virology 1:190–206.

Masson, F., Calderon Copete, S., Schüpfer, F., Garcia-Arraez, G., and Lemaitre, B. (2018) *In vitro* culture of the insect endosymbiont *Spiroplasma poulsonii* highlights bacterial genes Iinvolved in host-symbiont interaction. mBio 9:e00024–18.

McCutcheon, J.P., Boyd, B.M., and Dale, C. (2019) The life of an insect endosymbiont from the cradle to the grave. Curr Biol 29:R485–R495.

McLean, A.H., van Asch, M., Ferrari, J., and Godfray, H.C. (2011) Effects of bacterial secondary symbionts on host plant use in pea aphids. Proc Biol Sci 278:760–766.

Moran, N.A., McCutcheon, J.P., and Nakabachi, A. (2008) Genomics and evolution of heritable bacterial symbionts. Annu Rev Genet 42:165–190.

Murooka, Y., and Harada, T. (1979) Expansion of the host range of coliphage P1 and gene transfer from enteric bacteria to other gram-negative bacteria. Appl Environ Microbiol 38:754–757.

Murphy, K.C., and Campellone K.G. (2003) Lambda Red-mediated recombinogenic engineering of enterohemorrhagic and enteropathogenic *E. coli*. BMC Mol Biol 4:11.

Nováková, E., and Hypsa, V. (2007) A new *Sodalis* lineage from bloodsucking fly *Craterina melbae* (Diptera, Hippoboscoidea) originated independently of the tsetse flies symbiont *Sodalis glossinidius*. FEMS Microbiol Lett 269:131–135.

O’Connor, K.A., and Zusman, D.R. (1983) Coliphage P1-mediated transduction of cloned DNA from *Escherichia coli* to *Myxococcus xanthus*: use for complementation and recombinational analyses. J Bacteriol 155:317–329.

Oliver, K.M., Russell, J.A., Moran, N.A., and Hunter, M.S. (2003) Facultative bacterial symbionts in aphids confer resistance to parasitic wasps. Proc Natl Acad Sci USA 100:1803–1807.

Ornellas, E.P., and Stocker, B.A. (1974) Relation of lipopolysaccharide character to P1 sensitivity in *Salmonella typhimurium*. Virology 60:491–502.

Osborn, M.J. (1968) Biochemical characterization of mutants of *Salmonella typhimurium* lacking glucosyl or galactosyl lipopolysaccharide transferases. Nature 217:957–960.

Pais, R., Lohs, C., Wu, Y., Wang, J., and Aksoy, S. (2008) The obligate mutualist *Wigglesworthia glossinidia* influences reproduction, digestion, and immunity processes of its host, the tsetse fly. Appl Environ Microbiol 74:5965–5974.

Peters, J.E., and Craig, N.L. (2011) Tn*7*: smarter than we thought. Nat Rev Mol Cell Biol 2:806–814.

Pontes, M.H., Dale, C. (2006) Culture and manipulation of insect facultative symbionts. Trends Microbiol 14:406–412.

Pontes, M.H., and Dale, C. (2011) Lambda red-mediated genetic modification of the insect endosymbiont *Sodalis glossinidius*. Appl Environ Microbiol 77:1918–1920.

Pontes, M.H., Smith, K.L., De Vooght, L., Van Den Abbeele, J., and Dale, C. (2011) Attenuation of the sensing capabilities of PhoQ in transition to obligate insect-bacterial association. PLoS Genet 7:e1002349.

Rosner, J.L. (1972) Formation, induction, and curing of bacteriophage P1 lysogens. Virology 48:679–689.

Rubin, E.J., Akerley, B.J., Novik, V.N., Lampe, D.J., Husson, R.N., and Mekalanos, J.J. (1999) *In vivo* transposition of mariner-based elements in enteric bacteria and mycobacteria. Proc Natl Acad Sci USA. 96:1645–1650.

Sabri, A., Leroy, P., Haubruge, E., Hance, T., Frère, I., Destain, J., and Thonart P. (2011) Isolation, pure culture and characterization of *Serratia symbiotica* sp. nov., the R-type of secondary endosymbiont of the black bean aphid *Aphis fabae*. Int J Syst Evol Microbiol 61:2081–2088.

Santos-Garcia, D., Silva, F.J., Morin, S., Dettner, K., and Kuechler, S.M. (2017) The all-rounder *Sodalis*: a new bacteriome-associated endosymbiont of the lygaeoid bug *Henestaris halophilus* (Heteroptera: Henestarinae) and a critical examination of its evolution. Genome Biol Evol 9:2893–2910.

Schlechter, R.O., Jun, H., Bernach, M., Oso, S., Boyd, E., Muñoz-Lintz, D.A., et al. (2018) Chromatic bacteria - A broad host-range plasmid and chromosomal insertion toolbox for fluorescent protein expression in bacteria. Front Microbiol 9:3052.

Silipo, A., and Molinaro, A. (2010) The diversity of the core oligosaccharide in lipopolysaccharides. Subcell Biochem 53:69–99.

Smith, W.A., Oakeson, K.F., Johnson, K.P., Reed, D.L., Carter, T., Smith, K.L., et al. (2013). Phylogenetic analysis of symbionts in feather-feeding lice of the genus *Columbicola*: evidence for repeated symbiont replacements. BMC Evol Biol 13:109.

Šochová, E., Husník, F., Nováková, E., Halajian, A., and Hypša, V. (2017) *Arsenophonus* and *Sodalis* replacements shape evolution of symbiosis in louse flies. PeerJ 5:e4099.

Soucy, S.M., Huang, J., Gogarten, J.P. (2015) Horizontal gene transfer: building the web of life. Nat Rev Genet 16:472–482.

Sternberg, N., and Coulby, J. (1987a) Recognition and cleavage of the bacteriophage P1 packaging site (*pac*). I. Differential processing of the cleaved ends *in vivo*. J Mol Biol 194:453–468.

Sternberg, N., and Coulby J. (1987b) Recognition and cleavage of the bacteriophage P1 packaging site (*pac*). II. Functional limits of *pac* and location of *pac* cleavage termini. J Mol Biol 194:469–479.

Streicher, S., Gurney, E., and Valentine, R.C. (1971) Transduction of the nitrogen-fixation genes in *Klebsiella pneumoniae*. Proc Natl Acad Sci USA. 68:1174–1177.

Thomason, L.C., Costantino, N., and Court, D.L. (2007) *E. coli* genome manipulation by P1 transduction. Curr Protoc Mol Biol Chapter 1:Unit 1.17.

Thomason, L.C., Sawitzke, J.A., Li, X., Costantino, N., and Court, D.L. (2014) Recombineering: genetic engineering in bacteria using homologous recombination. Curr Protoc Mol Biol 106:1.16.1–39.

Toh, H., Weiss, B.L., Perkin, S.A., Yamashita, A., Oshima, K., Hattori, M., and Aksoy, S. (2006) Massive genome erosion and functional adaptations provide insights into the symbiotic lifestyle of *Sodalis glossinidius* in the tsetse host. Genome Res 16:149–156.

Touchon, M., Moura de Sousa, J.A., and Rocha, E.P. (2017) Embracing the enemy: the diversification of microbial gene repertoires by phage-mediated horizontal gene transfer. Curr Opin Microbiol 38:66–73.

Westwater, C., Schofield, D.A., Schmidt, M.G., Norris, J.S., and Dolan, J.W. (2002) Development of a P1 phagemid system for the delivery of DNA into Gram-negative bacteria. Microbiology 148:943–950.

Wilkinson, R.G., and Stocker, B.A. (1968) Genetics and cultural properties of mutants of *Salmonella typhimurium* lacking glucosyl or galactosyl lipopolysaccharide transferases. Nature 217:955–957.

Wilkinson, T.L. (1998) The elimination of intracellular microorganisms from insects: an analysis of antibiotic-treatment in the pea aphid (*Acyrthosiphon pisum*). Comp Biochem Physiol Ser A119:871–881.

Wolf-Watz, H., Portnoy, D.A., Bölin, I., and Falkow, S. (1985) Transfer of the virulence plasmid of *Yersinia pestis* to *Yersinia pseudotuberculosis*. Infect Immun 48:241–253.

Yarmolinsky, M.B., Sternberg, N. (1988) Bacteriophage P1. In The Bacteriophages. Calendar, R. (ed). New York: Plenum Press, vol 1. pp. 291–438.

Zaidman-Rémy, A., Vigneron, A., Weiss, B.L., and Heddi, A. (2018) What can a weevil teach a fly, and reciprocally? Interaction of host immune systems with endosymbionts in *Glossina* and *Sitophilus*. BMC Microbiol 18(Suppl 1):150.

